# A combined meta-barcoding and shotgun metagenomic analysis of spontaneous wine fermentation

**DOI:** 10.1101/098061

**Authors:** Peter R. Sternes, Danna Lee, Dariusz R. Kutyna, Anthony R. Borneman

**Affiliations:** The Australian Wine Research Institute, PO Box 197, Glen Osmond, South Australia, 5064; Department of Genetics and Evolution, University of Adelaide, South Australia. Australia. 5000

## Abstract

Wine is a complex beverage, comprising hundreds of metabolites produced through the action of yeasts and bacteria in fermenting grape must. To ensure a robust and reliable fermentation, most commercial wines are produced via inoculation with commercial strains of the major wine yeast, *Saccharomyces cerevisiae*. However, there is a growing trend towards the use of uninoculated or “wild” fermentations, in which the yeasts and bacteria that are naturally associated with the vineyard and winery, perform the fermentation. In doing so, the varied metabolic contributions of the numerous non-*Saccharomyces* species in this microbial community are thought to impart complexity and desirable taste and aroma attributes to wild ferments in comparison to their inoculated counterparts.

In order the map the microflora of spontaneous fermentation, metagenomic techniques were used to characterize and monitor the progression of fungal species in several wild fermentations. Both amplicon-based ITS phylotyping (meta-barcoding) and shotgun metagenomics were used to assess community structure. While providing a sensitive and highly accurate means of characterizing the wine microbiome, the shotgun metagenomic data also uncovered a significant over-abundance bias in the ITS phylotyping abundance estimations for the common non-*Saccharomyces* wine yeast genus *Metschnikowia*.

## INTRODUCTION

Wine is a complex beverage, comprising thousands of metabolites that are produced through the action of yeasts and bacteria in fermenting grape must. When grapes are crushed and allowed to ferment naturally, a complex microbial succession of yeasts and bacteria is generally observed. In the very early stages of fermentation, aerobic and apiculate yeasts, and yeast-like fungi from genera such as *Aureobasidium*, *Rhodotorula*, *Pichia*, *Candida*, *Hanseniaspora* and *Metschnikowia*, which reside on the surface of intact grape berries or winery equipment, represent the majority of the microbiota (1).

However most of these species, especially the aerobic yeasts, succumb early in the succession of the fermentation in response to falling oxygen levels and increasing ethanol. Mildly fermentative yeasts, such as *Hanseniaspora uvarum*, *Candida stellata*, *Metschnikowia pulcherrima*, *Torulaspora delbrueckii* and *Lachancea thermotolerans* can proliferate and survive well into the fermentation, but fall in numbers as ethanol levels increase, although it has been reported that *C*. *stellata* can survive up to 12% ethanol and complete fermentation (2–5).

Despite the vastly higher numbers of non-*Saccharomyces* yeasts early in the fermentation process, the major wine yeast, *Saccharomyces cerevisiae* is responsible for the bulk of the ethanolic fermentation. However *S*. *cerevisiae* is not readily isolated from intact grape berries and is therefore generally found in very low numbers at the start of fermentation (6, 7). Regardless, due to its higher fermentative ability, growth rate and tolerance to ethanol, *S*. *cerevisiae* supplants the various non-*Saccharomcyes* yeasts, becoming the dominant species from mid-fermentation such that an almost monoculture of this one species is established by the end of fermentation.

While traditional microbiological techniques have provided important insights into microbial succession that occurs in spontaneous ferments, both the breadth of ferments investigated and the depth at which individual species contributions could be resolved has been limited. Recent advances in culture-independent methods for species analysis, such as amplicon based phylotyping (also known as meta-barcoding) and metagenomics provide a high-throughput means to analyze large numbers of microbiological samples at great depth (8). Accordingly, these techniques are now being adapted for the study of wine fermentation, with several amplicon based methods, being used to investigate vineyard and wine microbiomes (9–13).

However, despite many studies using amplicon phylotyping techniques, there are still concerns regarding biases that may be inherent in the process, due to uneven PCR amplification or unequal copy number of the ribosomal repeat (9, 14). In order to address some of these limitations, metagenomic techniques are being used to determine species abundance from shotgun sequencing of mixed samples. These techniques generally rely on read mapping, either to collections of curated marker genes or whole genomes, making them reliant on reference sequences that are available (15). As yet, shotgun metagenomics has not been applied to the study of wine fermentation, or to assess the accuracy of amplicon based abundance estimates of wine fermentation.

In order to address this, fungi-specific ITS-phylotyping was performed over four key fermentation stages in five independent commercial Chardonnay juice fermentations in triplicate. Full shotgun metagenomic sequencing was also performed for twenty of these samples. Comparison of the ITS-phylotyping and shotgun data, uncovered a major amplicon bias that existed for the genus *Metschnikowia*, providing the means to normalize other ITS-datasets in which this species is abundant.

## MATERIALS AND METHODS

### Laboratory ferments

For each of the laboratory-scale wild ferments, 20 L of Chardonnay grape juice was obtained directly from winery fermentation tanks immediately after crushing. Each 20 L sample was then split into three separate 3 L glass fermenters fitted with air-locks and fermented at 20 °C with daily stirring.

Samples were taken daily for measurement of Baumé (Bé) and 50 mL samples taken at the time of transfer (D0), after ~1 Bé reduction in sugar concentration (D1), 50 % sugar reduction (D2), and then at dryness (0-3 Bé, D3) for enumeration by both microbiological plating and ITS/metagenomic analysis.

For classical microbiological plating, serial dilutions were plated onto both WL nutrient agar (Oxoid) for estimation of total yeast numbers and lysine medium (Oxoid) for estimation of total non-*Saccharomyces* yeasts.

### DNA preparation

For each sample, 50 mL of fermenting juice was centrifuged for 10 mins at 10,000 g, washed in 20 mL PBS, re-centrifuged and then frozen at −80 °C until processed. Total DNA was extracted from washed must pellets using the PowerFood Microbial DNA Isolation Kit (Mobio).

### ITS-amplicon preparation and anlaysis

Analysis of ITS abundance from ferment samples was performed using two-step PCR amplification followed by next-generation amplicon sequencing (Fig. S1). First-round amplification of the ITS region was performed using the fungal-specific primers BITS (ACCTGCGGARGGATCA) and B58S3 (GAGATCCRTTGYTRAAAGTT) (9) which were modified to include both an inline barcode and Illumina adaptor sequences BITS-F1-N*xxx* and BITS-R1-N*xxx* (Table 1). One nanogram of DNA was used in each first-round PCR (20-30 cycles, 55 °C annealing, 30 sec extension, KAPA 2G Robust polymerase). Second-round amplification was performed using the Illumina adaptor sequence present in the first-round primers as an amplification target, with the remaining sequences required for dual-indexed sequencing on the Miseq platform added via overhang PCR (Fig. S1). For each sample, 2 uL of first-round PCR product was used (15 cycles, 55 °C annealing, 30 sec extension, KAPA 2G Robust polymerase). Following PCR, all samples were mixed into a single batch and column purified (minEulte, Qiagen). ITS-amplicon pools were sequenced on the Illumina Miseq sequencing platform using 2 x 300 bp paired-end chemistry (Ramaciotti Centre for Functional Genomics, Australia).

**Table 1.**
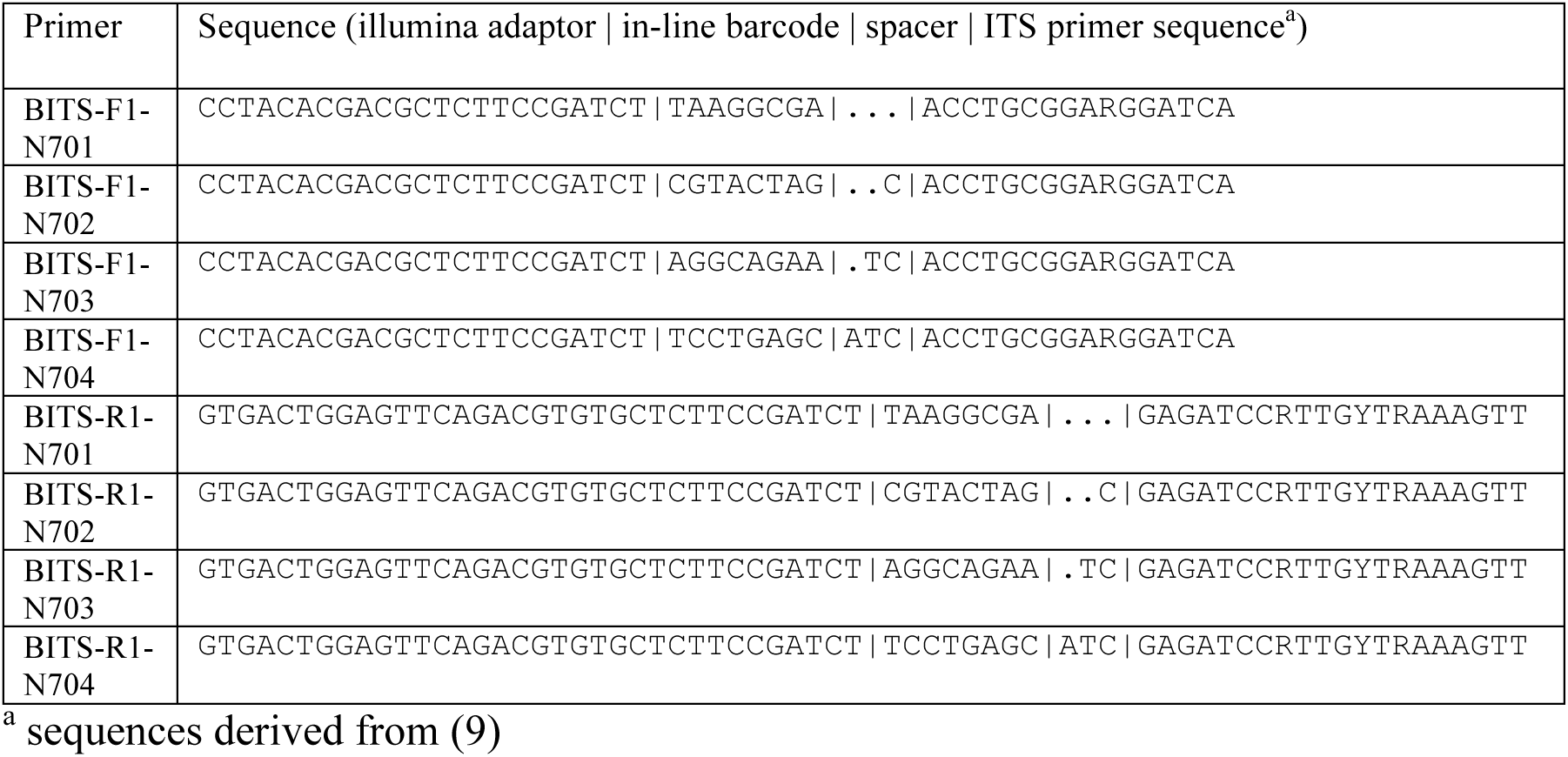
ITS amplification primers used in this study.

Following sequencing, raw sequence data were quality trimmed (Trimmomatic v0.22 (16); TRAILING:20 MINLEN:50), adaptor trimmed at the 3′ end to remove ITS adaptor sequences (cutadapt 1.2.1 (17)) and the individual read pairs were overlapped to form single synthetic reads (FLASH 1.2.11 (18); --max-overlap 1000 --allow-outies). These synthetic reads were then trimmed at both the 5′ and 3′ ends to remove any remaining Illumina adaptors that were directly adjacent to the inline barcodes (cutadapt 1.2.1 (17); -a AGATCGGAAG -g CTTCCGATCT -e 0.1 -overlap 6) and sequentially partitioned according to the specific combination of inline barcode sequences at both the 5′ and 3′ end of each synthetic read using FASTX-Toolkit (v0.0.13; fastx_barcode_splitter.pl --bol--mismatches 1;http://hannonlab.cshl.edu/fastx_toolkit/).

In order to calculate the abundance of individual amplicons the entire dataset for all of the samples were dereplicated with USEARCH (v7.0.1990; -derep_full length, -size-out; (19)) and each dereplicated OTU renamed according to the md5 checksum of the OTU sequence to provide unambiguous comparison of identical OTUs across experiments. Dereplicated OTUs were then clustered using SWARM (-z --differences 1 --fastidious (20)), with a minimum final OTU size of 10 implemented using custom scripts.

Following clustering, the likely taxonomic identity of the representative sequence of each OTU was determined using the assign_taxonomy.py module of QIIME using a modified form of the standard QIIME UNITE database in which any unclassified or unidentified sequences were removed and each ITS region was trimmed to the extent of the BITS primers used for the original ITS amplification (-t sh_taxonomy_qiime_ver6_dynamic_s_10.09.2014.txt -r sh_refs_qiime_ver6_dynamic_s_10.09.2014.BIT.unclassified.unidentified.fasta -m uclust --uclust_similarity = 0.98 --uclust_max_accepts=10 --uclust_min_consensus_fraction = 0.4; (21)). In addition to the edited UNITE database, OTU annotations were also performed with an augmented version of the database in which several wine-specific, manually-curated reference sequences were added and three UNITE reference sequences that were found to have erroneous annotations were either edited or removed (Supplemental File 1).

Once the results were established for the full dataset, individual dereplicated OTUs from each sample were matched back to those of the full dataset using custom scripts to provide a directly comparable and standardized assignment of each individual experimental result within the overall dataset. Final results were assembled in QIIME tabular format using custom scripts (Table S2).

Multidimensional data analysis was performed with the R phyloseq package (22) using principal coordinate analysis (PCoA) and Bray Curtis dissimilarity measures based upon the 30 most abundance OTUs across the samples.

### Shotgun metagenomics analysis

DNA from four control populations, 16 fermentation samples and two winery samples were subjected to whole genome metagenomic sequencing. Random sequencing libraries were prepared using the Truseq nano protocol (Illumina) with a ~350 bp insert size. Sequencing libraries were then pooled and run across three lanes of Illumina Hiseq 2 x 100 bp chemistry (Ramaciotti Centre for Functional Genomics, Australia).

Following sequencing, raw sequence data was first filtered to limit contaminating grapevine sequences by aligning each set of sequences against the Pinot Noir grapevine genome (CAAP00000000.3; (23)) using Bowtie2 v2.2.5 in unpaired mode (24). All unaligned reads for which both reads in a pair failed to align to the grapevine genome were retained for further analysis.

In order to provide a reference sequence for read mapping, whole genome sequences were collected, where possible, from a combination of species comprising either known grape and wine microbiota (including bacteria) or other fungal species identified as being present in the fermentations analyzed in this study via ITS-phylotyping. (Table S3). This reference sequence was divided up into discrete windows of 10 kb using Bedtools2 (v2.24.0; makewindows -w 10000; (25)).

Each of the filtered shotgun datasets were then aligned to this reference set using Bowtie2 in paired-end mode with unaligned reads saved for later analysis (--fr --maxins 1500 --no-disconcordant --no-unal --un-conc (24)). The resultant .sam files were sorted and converted to .bam format and filtered for low-quality alignments using Samtools (v1.2; view -bS -q 10 | sort; (26)). For each .bam file the total read coverage in each 10 kb reference window was calculated using Bedtools2 (v2.24.0; coverage -counts; (25)), with the mean, median and adjusted mean (retain mean if >= 20 % of the windows in that species contained >= 1 read, otherwise mean value of 0 applied) calculated from the bed window values for each species in each sample using custom scripts. In addition to coverage values, the average identity of each mapped read was calculated for each window using custom scripts that counted the number of mismatches per read (Bowtie2 XM: tag for each read) compared to overall read length.

### *De novo* metagenomic assembly

For the assembly of uncultivated sequences that were unrepresented in early versions of the shotgun reference collection, reads that failed to align during the shotgun metagenomic analysis were *de novo* assembled using SPADES (v3.5.0; --sc --careful; (27)). The likely taxonomic source of each contig was estimated using BLASTX (ncbi_blast-2.2.31+; -task blastx-fast -outfmt "7 std sscinames" -max_target_seqs 20) against the non-redundant database (nr; date 02/14/2015) and extracting the taxonomic source of the best blast hit. Contigs were then partitioned according this taxonomic grouping at the genus level, with genera being manually combined where appropriate.

### Data availability

All sequencing data, including ITS barcoding and shotgun metagenomic sequencing have been deposited in Genbank under the Bioproject accession number PRJNA305659.

## RESULTS AND DISCUSSION

### Analysis of microbial communities in wild ferments

In order to study the reproducibility and applicability of laboratory-scale uninoculated ferments, five Chardonnay grape musts (Y1, Y2, Y3, T1 and T2), which were each destined to undergo winery-scale uninoculated fermentation (sourced from two different wineries), were fermented at laboratory scale in triplicate (Table 2). Fermenting musts were tracked for sugar consumption via refractometry, with samples taken for analysis at four key time points (D0, at inoculation; D1, after 1 Baumé (Bé) drop; D2, ~ 6 Bé; D3, ~3 Bé). According to selective plating, all of the ferments showed classical microbiological progression, with *non-Saccharomyces* species showing an initial increase in numbers, followed by steady decline while *Saccharomyces spp.* greatly increased in number before reaching a plateau late in ferment. All ferments proceeded to dryness, with sample Y1 being the fastest (12 days) and sample T1 taking the longest time (27 days).

**Table 2.** Fermentation samples used in this study

| Sample | Source | Grape Variety | Vineyard/Winery Location | Matching LAB sample |
| --- | --- | --- | --- | --- |
| T1 | LAB | Chardonnay | Adelaide Hills, SA/Barossa Valley, SA |  |
| T2 | LAB | Chardonnay | Adelaide Hills, SA/Barossa Valley, SA |  |
| Y1 | LAB | Chardonnay | Eden Valley, SA/Barossa Valley SA |  |
| Y2 | LAB | Chardonnay | Eden Valley, SA/Barossa Valley SA |  |
| Y3 | LAB | Chardonnay | Adelaide Hills, SA/Barossa Valley, SA |  |

### Species abundance estimation via ITS amplicon analysis

A total of 66 samples were analyzed comprising control populations (n=6) and laboratory scale fermentations (n=60) (Table S1). DNA was isolated from the pelleted fraction of each must sample, with a two-step PCR performed using sequences designed to amplify the fungal ITS region (9), while adding experiment-specific inline barcodes and appropriate adaptors for sequencing on the Illumina sequencing platform (Fig. S1). Following sequencing and barcode and adaptor trimming, 8.8 million reads were assigned across the samples (Table S1), with an average of over 100,000 reads per sample.

In order to consistently describe and compare the number of OTUs across the samples, all 8.8 million reads were first analyzed as a large single batch. Dereplication (19), OTU clustering (20) and taxonomic assignment (21), of this combined dataset resulted in the production of a single OTU table that encompassed all the of OTUs from across all 78 samples. Abundance measurements of each individual dereplicated OTU from each sample were then mapped to this combined data table to derive the contribution of each experiment to the collective data set (Table S2).

### Control populations

Given previous concerns regarding the accuracy of ITS-amplicon profiling (9), two different control populations were assembled, in triplicate, from individual cultures of eight common wine-associated yeasts, representing seven different species and six different genera (Table 3). By comparing the results of the ITS-amplicon profiling of these samples with those expected from estimated numbers of input cells, nearly all species estimates were within two-fold of their expected value, despite cell concentrations differing across five orders-of-magnitude (Table 3). However, the results for *Metschnikowia* appeared to be reproducibly over estimated in both control populations (18.6 and 10.5 fold), indicating that this species may display significant amplicon bias for the ITS region relative to the other samples used in the control populations.

**Table 3.** Control populations

| Strain | Species | Control mix 1 | Total OTUs | ITS abundance<sup>a</sup> (ratio) | Shotgun abundance<sup>a</sup> (ratio) | Control mix 2 | Total OTUs | ITS abundance<sup>a</sup> (ratio) | Shotgun abundance<sup>a</sup> (ratio) |
| --- | --- | --- | --- | --- | --- | --- | --- | --- | --- |
| AWRI796 | *Saccharomyces cerevisiae* | 1x10<sup>6</sup> | 3 | 1 x10<sup>6</sup> (1) | 1 x10<sup>6</sup> (1) | 1x10<sup>8</sup> | 11 | 2 x10<sup>8</sup> (1) | 2 x10<sup>8</sup> (1) |
| AWRI1498 | *Saccharomyces cerevisiae* | 1x10<sup>4</sup> |  |  |  | 1x10<sup>8</sup> |  |  |  |
| AWRI1149 | *Metschnikowia pulcherrima* | 1x10<sup>4</sup> | 7 | 1.9 x10<sup>5</sup> (18.6) | 8.8 x 10<sup>3</sup> (0.9) | 1x10<sup>6</sup> | 7 | 1.1x10<sup>7</sup> (10.5) | 6.5 x 10<sup>5</sup> (0.7) |
| AWRI1152 | *Torulaspora delbrueckii* | 1x10<sup>6</sup> | 1 | 7.3 x10<sup>5</sup> (0.7) | 4.8 x 10<sup>5</sup> (0.5) | 1x10<sup>5</sup> | 1 | 6.6 x 10<sup>4</sup> (0.7) | 4.7 x 10<sup>4</sup> (0.5) |
| AWRI1157 | *Debaryomyces hansenii* | 1x10<sup>7</sup> | 1 | 8.4 x10<sup>6</sup> (0.8) | 2.9 x 10<sup>6</sup> (0.3) | 1x10<sup>3</sup> | 1 | 2.4 x 10<sup>3</sup> (2.4) | 0.0 |
| AWRI1176 | *Saccharomyces uvarum* | 1x10<sup>3</sup> | 1 | 5.7 x10<sup>2</sup> (0.6) | 1.8 x 10<sup>3</sup> (1.8) | 1x10<sup>5</sup> | 3 | 7.9 x 10<sup>4</sup> (0.8) | 1.1 x 10<sup>5</sup> (1.1) |
| AWRI1274 | *Haneniaspora uvarum* | 1x10<sup>8</sup> | 4 | 1.1 x 10<sup>8</sup> (1.1) | 1.3 x 10<sup>8</sup> (1.3) | 1x10<sup>4</sup> | 2 | 6.1 x 10<sup>4</sup> (6.1) | 4.4 x 10<sup>4</sup> (4.4) |

<sup>a</sup> normalised using the abundance of *S. cerevisiae*

### Uninoculated ferments

ITS-amplicon analysis of laboratory scale wild ferments showed that there was a high degree of reproducibility between each of the three biological triplicates (r^2^ 0.95 ± 0.02; Fig. S2). All of the fermentations displayed an expected microbiological succession, beginning with a diverse and variable collection of fungi that progressively resolved into a population that was dominated by the major wine yeast *S*. *cerevisiae* (Fig. 1A). Multidimensional analysis (Bray-Curtis) of the ferments showed that while T2, Y1 an Y2 could be broadly classified as being dominated by *Metschnikowia* and *Hanseniaspora* at the D0 and D1 time points, the Y3 ferment was almost devoid of these genera, with the ferment characterized by high levels of *Aureobasidium* and *Rhodotorula*, primarily at D0 (Fig. 1B). T1, the slowest ferment, displayed a highly diverse D0 population of *Rhodotorula, Cladosporium* and *Aureobasidium,* which progressed through a *Hanseniaspora-dominated* phase at D1 and finally to *S. cerevisiae* at D2/D3.

**Figure 1.**
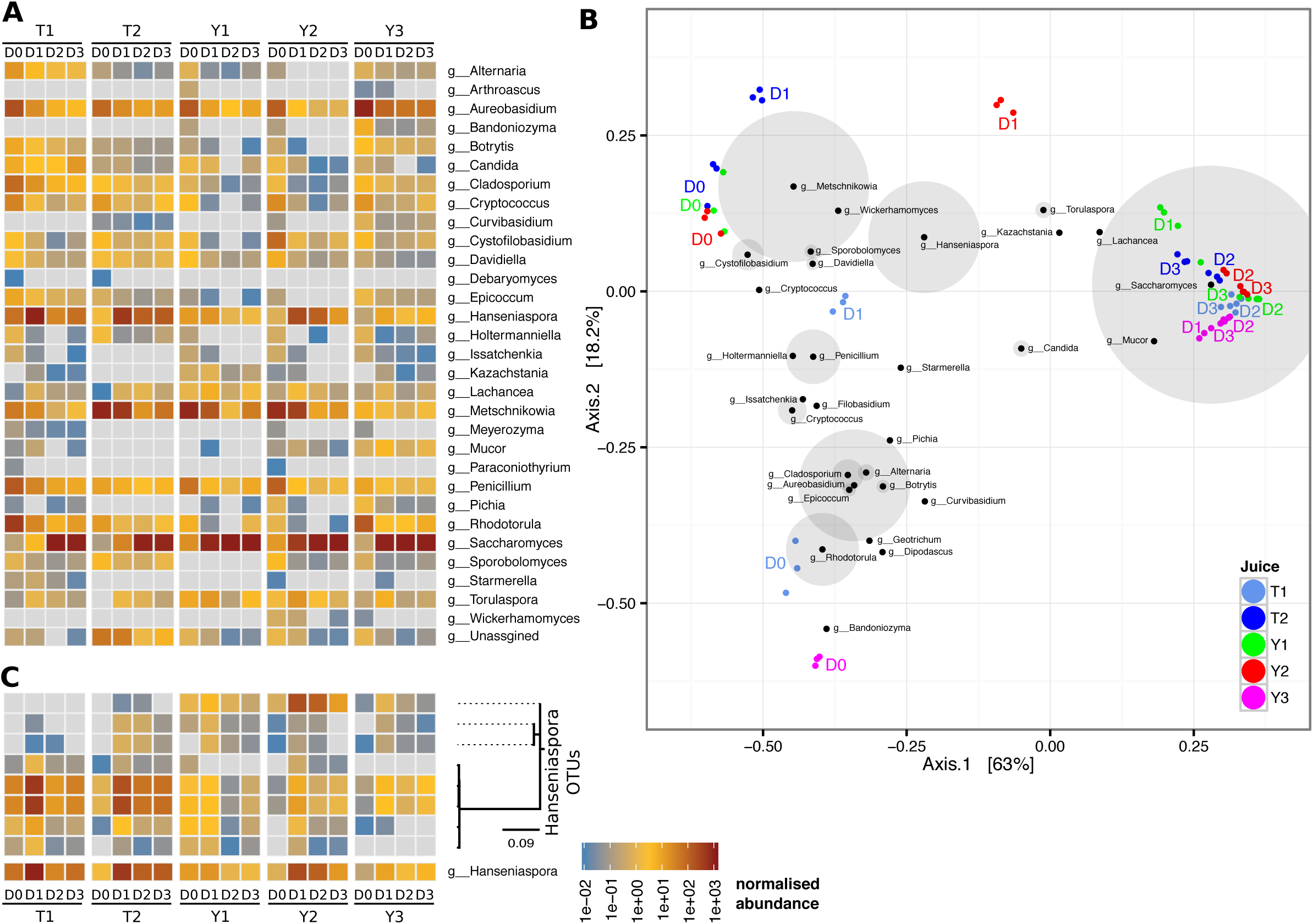
ITS amplicon abundance of uninoculated ferments. (A) Laboratory-scale ferments analyzing four fermentation time points in five different musts in triplicate. In both plots, ITS sequences are grouped by genus and are colored-coded by their normalized abundance (reads per thousand reads) (B) Dissimilarity analysis of ITS-amplicon abundance. Triplicate samples from each time point were subjected to Bray-Curtis dissimilarity analysis. The weightings of the top 30 genera are overlaid on the plot, with the size of the grey circles around each node proportional to the total abundance of each genus across all samples (no shading for nodes >5000 counts). (C) Species-level ITS assignment for the genus *Hanseniaspora.* The individual abundance measurements for the eight OTUs that comprise the g_ Hanseniaspora category are shown, grouped by phylogenetic distance. Abundance values are presented as in Fig 1A.

The use of the fungal ITS marker also allowed for species-level assignment of many OTUs and there were several genera for which more than one species was encountered. For example, the genus *Hanseniaspora* was represented by a total of eight OTUs that could be grouped into at least five main species (by ITS sequence similarity) (Fig 1C). (*H*. *uvarum*, *H*. *opunitae*, *H*. *osmophila*, *H*. *vineae* and *H*. *guilliermondii*) with two species, *H*. *uvarum* and *H*. *opuntiae* having two distinct OTUs representing each species, but which displayed coordinated changes in abundance across both juice and time point. For these species, this argues that either there were multiple strains of each species present in the ferments (with slightly different ITS sequences) that were responding similarly, or that multiple OTU sequences were being produced per species (either due to heterogenous ITS repeats or PCR artefacts).

Interestingly, these two main categories of ferments that were observed (T2, Y1 and Y2 versus T1 and Y3) did not correlate with vineyard location, winery or the stage of vintage (Table 2). However, the overall difference in the location of the vineyards and wineries is relatively minor, with Eden Valley and the Adelaide Hills being geographically adjacent regions in South Australia. The driver of these striking differences in microbial starting populations and progressions therefore remains to be determined, however factors such as vineyard management and/or microclimate are likely to be involved (28–31).

### Shotgun metagenomics

While ITS-amplicon sequencing provided an in-depth analysis of variation across ferments, it has been widely accepted that the combination of ITS primer sequences, multiple rounds of PCR, and variation in the ITS repeat number can produce biases in the final abundance measurements (9, 14). In addition, unless a second primer set is employed, bacterial species are not covered by this analysis. In order to explore these potential biases in more detail and to potentially provide strain-level information, shotgun metagenomics, in which total DNA is extracted and directly sequenced, was employed on a total of twenty of the samples analyzed by ITS-amplicon sequencing (Table 4).

**Table 4.** Shotgun metagenomic alignment statistics

| Sample | Total reads | Alignment rate (%) |
| --- | --- | --- |
| Control1_A | 18,816,478 | 97.05 |
| Control1_C | 17,883,449 | 97.11 |
| Control2_A | 20,343,317 | 97.18 |
| Control2_C | 18,138,322 | 97.91 |
| T1D1_A | 20,027,063 | 88.15 |
| T1D1_B | 21,175,617 | 90.22 |
| T2D1_A | 18,173,778 | 90.12 |
| T2D1_B | 21,705,055 | 91.02 |
| T2D3_A | 21,282,134 | 97.12 |
| T2D3_B | 21,112,818 | 96.84 |
| Y1D1_A | 20,196,267 | 96.04 |
| Y1D1_B | 20,404,693 | 96.39 |
| Y2D1_A | 18,384,104 | 93.13 |
| Y2D1_B | 18,960,301 | 92.61 |
| Y2D3_A | 20,731,661 | 96.66 |
| Y2D3_B | 19,327,663 | 96.75 |
| Y3D1_A | 19,843,426 | 94.48 |
| Y3D1_B | 21,559,192 | 94.29 |
| Y3D3_A | 19,533,468 | 96.69 |
| Y3D3_B | 19,258,370 | 95.77 |

Given that reference genome sequences exist for many wine associated microbes, a mapping abundance strategy was used to analyze the shotgun data. A representative collection of reference genomes was therefore assembled from existing genomic resources for fungal and bacterial genera that were known, or suspected of being wine-associated (Table S3). However, attempts at aligning to a preliminary reference genome set for shotgun abundance estimation (see below), resulted in up to 15% of reads being unable to be aligned. In order to determine if this was due to a lack of suitable reference genomes for key species that were represented in the shotgun data, all unaligned sequences were subsequently *de novo* assembled, and the resulting contigs partitioned according to their likely genus. Four genera were represented by at least 500 kb of sequence, although three of these (*Rhodosporidium*, *Rhodotorula* and *Microbotryum*) were combined and ascribed to a single major assembly product, *Rhodosporidium* (n=1539, 17.7 Mb). This was despite three other species of *Rhodosporidium* and *Rhodotorula* being present in the reference dataset, such that these contigs are likely to represent the entire genome of an additional species within this genus. Likewise, the fourth group of contigs (n=76, 644 kb) was ascribed to *Aureobasidium spp,* despite the presence of a reference sequence for *Aureobasidium pullulans*. However, in this instance, the strong correlation in abundance values obtained across the samples for these two different sequences point to these *de novo* contigs representing regions that are not conserved in the existing *A*. *pullulans* reference sequences, as the bulk of the reads were able to be aligned to the reference strain (Fig. S3). These sequences were subsequently added to the reference set for use in the shotgun abundance estimation.

### Estimating species abundance using shotgun metagenomic sequencing

The final reference genome set comprised 851 Mb of DNA that represented a total of 51 species (45 eukaryotic, 6 prokaryotic; Table S3). Alignment of each sample to the reference genome set resulted in the majority of reads being represented in the reference consortium, although this was highly sample dependent, with between 2 % and 11 % of reads not able to be matched to the reference genome set (Table 4). These sequences likely represent species for which an adequate reference genome did not exist and were present at too low of an abundance to produce an adequate *de novo* assembly from the unaligned pool. In order to determine the potential taxonomic source of these unaligned reads, the marker-gene metagenomic classifier MetaPhlAn (32) was used to classify the remaining reads from this dataset (Table S4). This indicated 40 % of the remaining reads to be of bacterial origin, 46 % from *Ascomycetous* fungi and 13 % viral. The most highly represented bacterial genera included *Acetobacter* (25 % total bacterial reads), *Curtobacterium* (14 %) and *Lactobacillus* (18 %). From the *Ascomycete spp*, half were predicted to be from *S*. *cerevisiae*, and likely represent mitochondrial reads (the mitochondrion was excluded from the reference genome set due to its variable copy number), with another 30% predicted to derive from an unclassified member of the family *Debaryomycetaceae*.

For those reads that were able to be matched to the reference set, estimations of species abundance were made from average read coverage values from discrete 10 kb windows across each genome (Fig. 2A). In addition to read depth, the average identity between each read and the reference to which it mapped was also recorded. This provided an estimate of the evolutionary distance between the particular genomic reference and the strains or species present in each sample. These identity values were generally above 99 % for the reference genomes, but were found to be significantly lower for reference sequences including *Mucor circinelloides*, *Pseudomonas syringae* and *Hanseniaspora valbyensis*, suggesting that the actual species or strains present in the fermentation were significantly different to the reference used.

**Figure 2.**
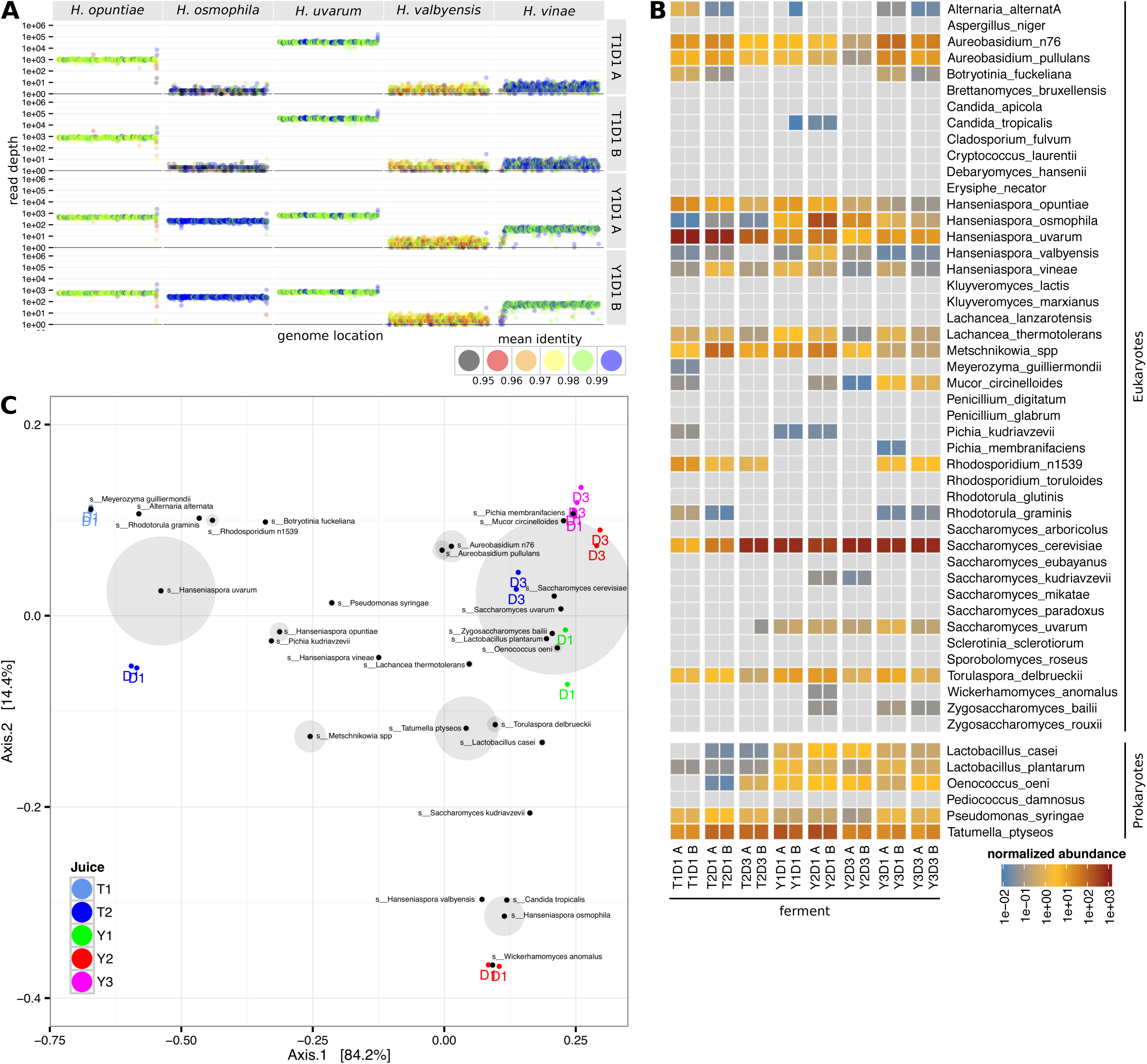
Shotgun metagenomic analysis of species. (A) Shotgun sequencing reads from each sample were mapped to the wine metagenome reference set. The total reads present in non-overlapping 10 kb windows across each genome were recorded relative to genomic location. In addition to total read number, the average identity of the reads in each window compared to the reference sequence was also calculated (id_factor). For clarity, only the abundance measures for species within the *Hanseniaspora* genus are depicted for two T2D1 and Y1D1 replicates. Results for all samples presented in Supp. Fig. R2). (B) Normalized average abundance values for each reference species in each sample. Values were normalized using total read numbers in each sample (including non-aligning reads) with final values represented per million reads in each 10 kb genomic window. (C) Bray-Curtis dissimilarity analysis of the shotgun abundance data. The weightings of each reference genome are overlaid on the plot, with the size of the grey circles around each node proportional to the total the abundance of each reference genome across all of the samples (no shading for nodes >10).

In order to provide single abundance values for each reference genome in each sample, overall abundance measurements were derived from the average read depth of all 10 kb windows in each genomic sequence. An additional filter of at least 20 % genome coverage was also applied to limit the effect of small numbers of windows with large coverage values, such as those derived from mis-mapping or potential small scale horizontal transfer events from very high abundance species against otherwise no-or low-abundance genomes (*e*.*g*. *S*. *cerevisiae* and *S*. *paradoxus*), from producing spurious abundance estimations. Using this technique, it was possible to detect the presence of 25 of the eukaryotic reference sequences and 5 prokaryotes, across five orders of magnitude (Fig. 2B). Comparing the shotgun metagenomic values obtained for the four control experiments, to those expected from the estimated numbers of input cells, showed that the outcomes of the shotgun analysis were within two-fold of each other in all but two cases, and highly correlated with the ITS results (R^2^ 0.99) (Table 3).

When ordinate analysis was used to compare the shotgun samples, the presence of high amounts of *Hanseniaspora spp*. in three D1 samples (T1, T2 and Y3) largely differentiated them from the two remaining samples. Within the three D1 samples that contained high levels of *Hanseniaspora spp*. the D1 T1 and T2 samples were primarily populated by *Hanseniaspora uvarum*, while the Y3 sample contained roughly equal proportions of *Hanseniaspora uvarum* and *Hanseniaspora osmophila* (Fig. 2C).

### Comparison of shotgun and ITS-amplicon data

In addition to comparing the control values, it was possible to extract values for comparison from the full shotgun and ITS-amplicon datasets by comparing the results from a total of 23 comparable taxonomic identifiers that were present in both experimental types (Fig. 3). For the majority of these taxonomic identifiers, the normalized abundance values recorded from the shotgun and ITS-amplicon experiments were highly correlated (R^2^ 0.93) and differed by two-fold or less, across a dynamic range of more than four orders of magnitude, with accuracy diminishing at levels below 100 reads/fragments per million. For those high abundance species that were not within a five-fold range, the previously identified ITS-amplicon over-estimation bias for *Metschnikowia spp*. was recapitulated, confirming that this is a bias inherent in ITS analysis for this species.

**Figure 3.**
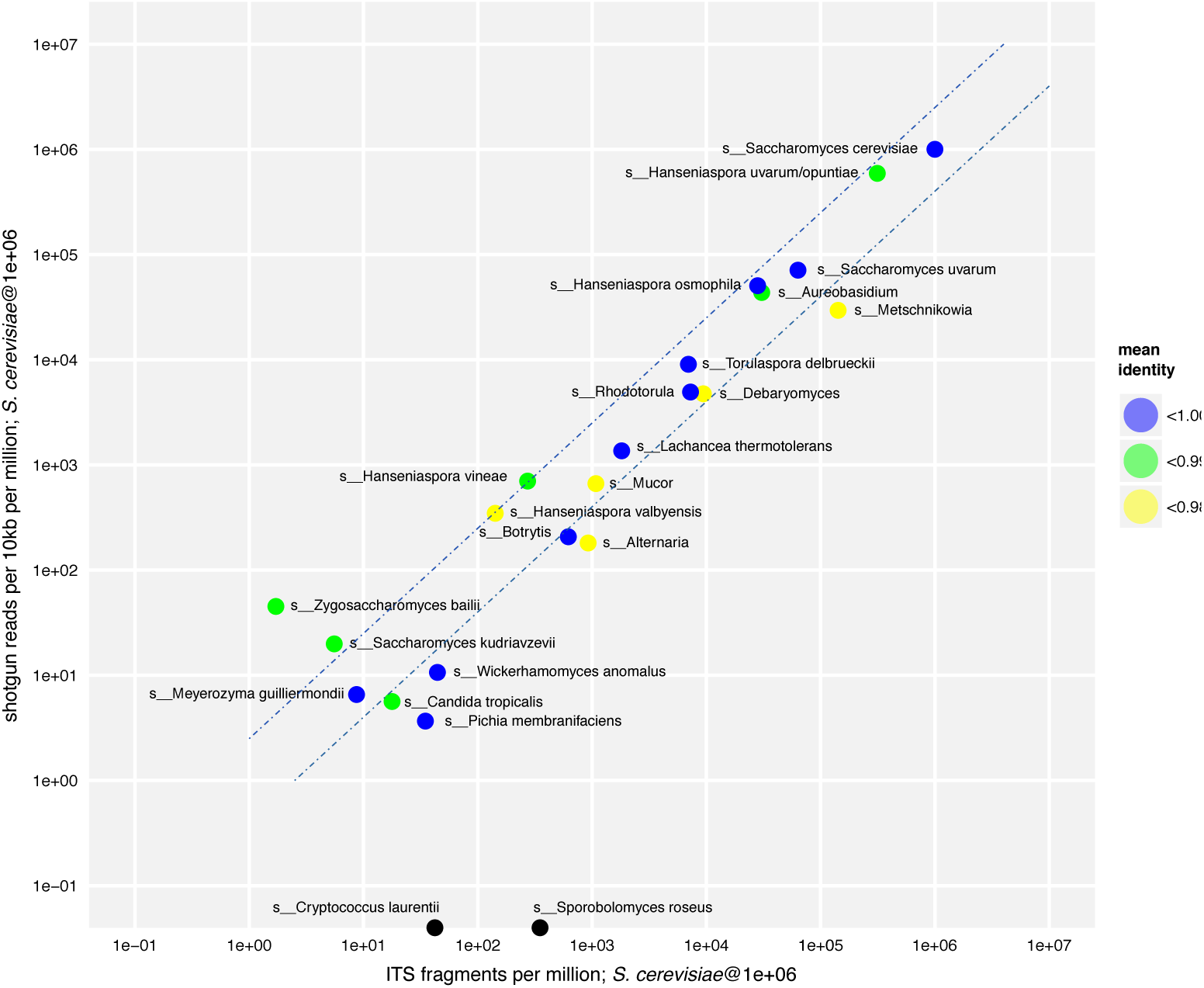
Comparison of ITS and shotgun abundance measurements. Normalized and abundance measurements were scaled for both the shotgun and ITS experimental designs relative to an abundance of *S*. *cerevisiae* of 1 million reads per million. Dashed lines represent two-fold variation between samples. The mean identity of the shotgun data relative to the reference genome used is also shown.

## Conclusions

Uninoculated wine ferments represent a complex and dynamic microbial community. Metagenomics and phylotyping are now allowing for the detailed analysis of large numbers of fermentation samples, shining a light on the composition of these microbial mixtures. While the ITS-phylotyping providing an accurate, high-throughput means to determine species abundance, the shotgun metagenomics uncovered at least one major example of amplicon bias, with the *Metschnikowia spp*. displaying a 10-fold over-representation. However, once biases such as these have been identified, they can be corrected in future ITS-phylotyping datasets to provide a more accurate species representation. As more shotgun metagenomic and single-strain *de novo* assemblies for key wine species become available, the accuracy of both ITS-amplicon and shotgun studies will greatly increase. This will provide a key methodology for deciphering the influence of the microbial community on the wine flavor and aroma and how winemaking interventions may be used to shape these outcomes.

## ACKNOWLEDGMENTS

Special Thanks to Louisa Rosa and Alana Seabrook of Yalumba and Alison Soden of Treasury Wine Estates for supplying must and wild fermentation samples and Paul Chambers for critical reading of this manuscript.

---

**Figure S1.**
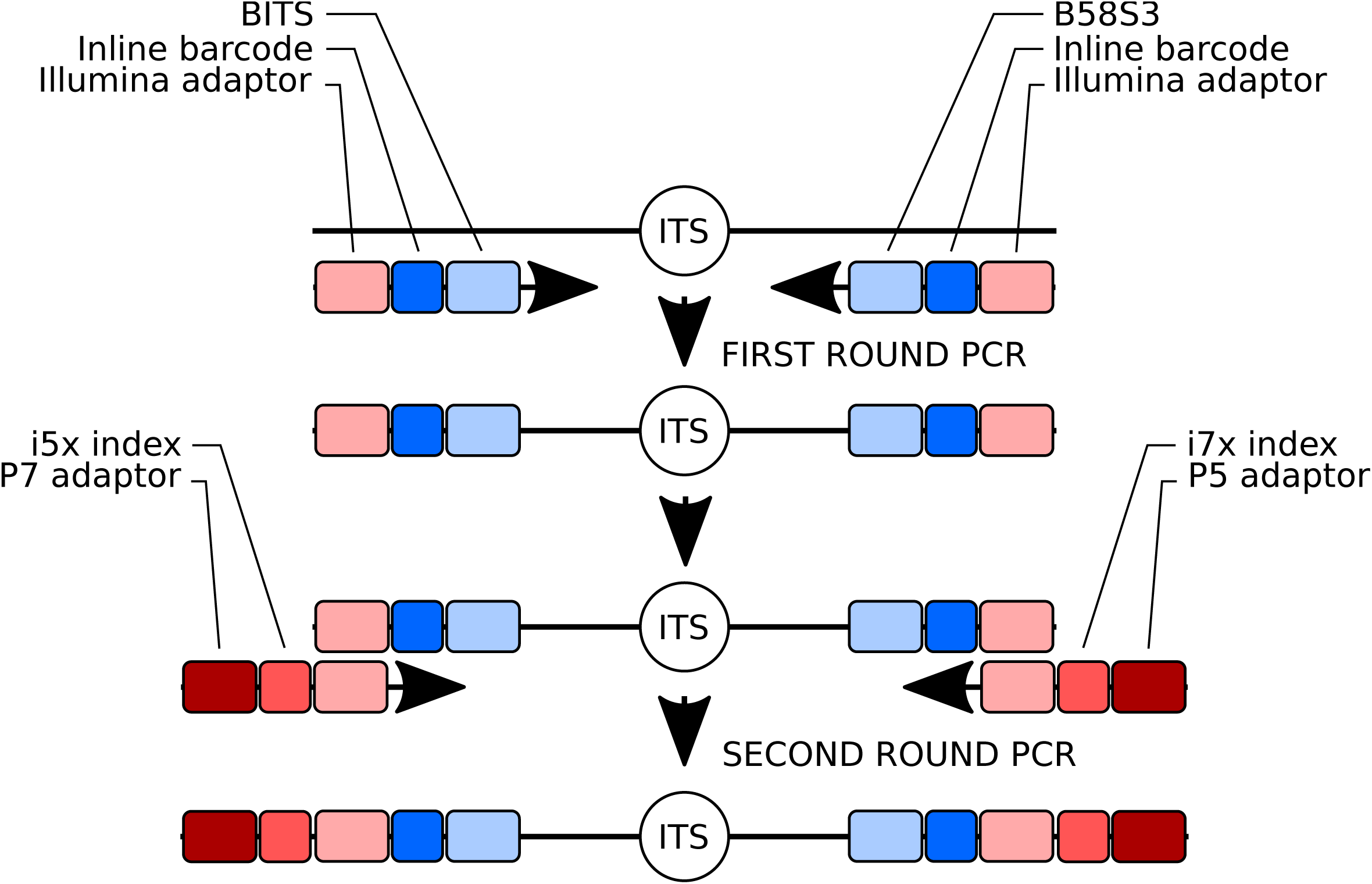
Two step amplification of the ITS region as an Illumina-ready amplicon. First round amplification uses BITS and B58S3 primers (9) fused to inline barcodes and adaptor sequences. Second round amplification takes advantage of the common Illumina adaptors to add Illumina indexing sequences and the P7 and P5 adaptors that are required for flow cell adherence and amplification.

**Figure S2.**
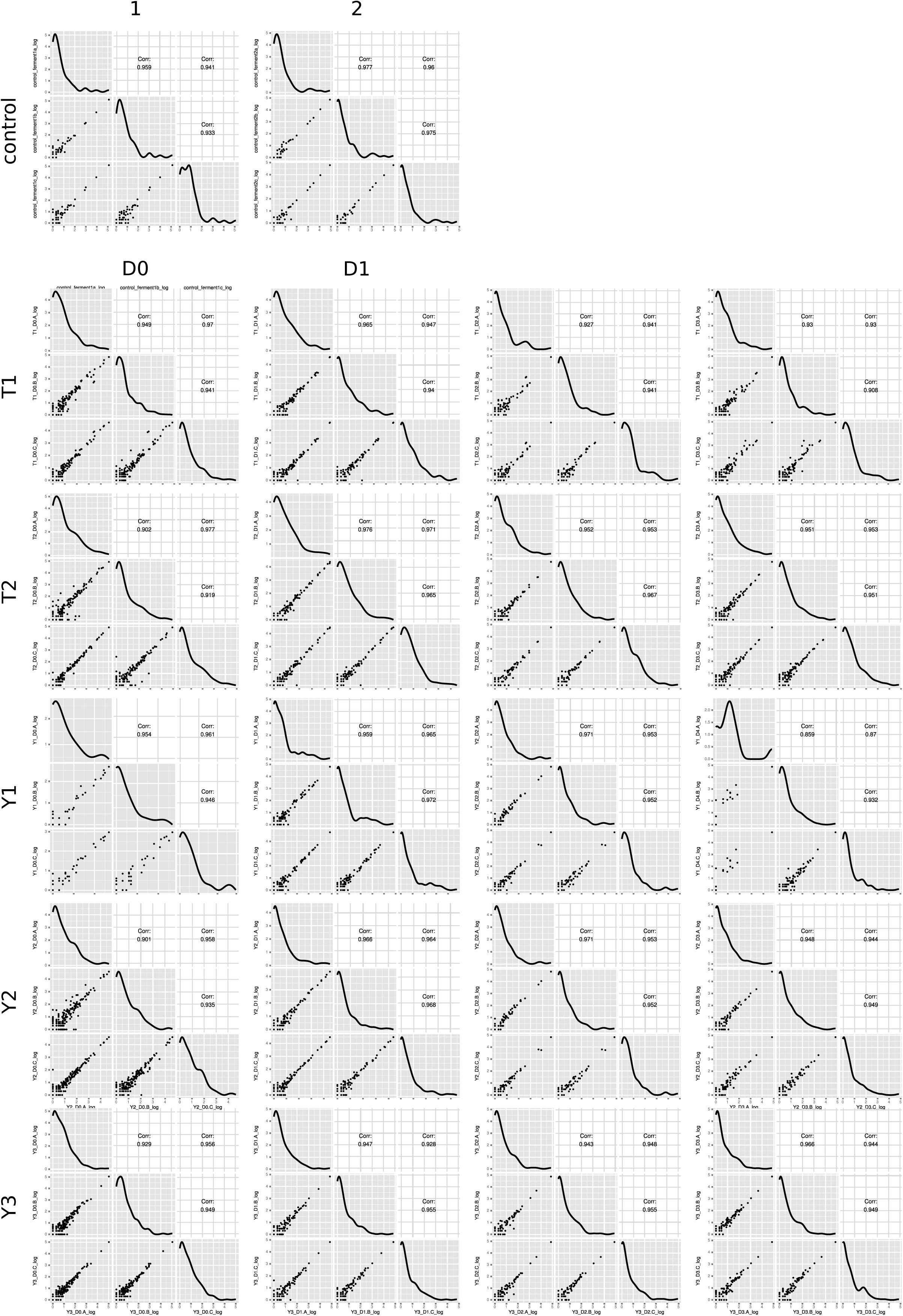
Correlation analysis of replicate ferments. Pair wise comparisons were performed between triplicate samples from the four fermentation time points across the five different juices. Raw abundance measurements were compared for all significant OTUs (>10 reads). R^2^ values are presented for each pairwise comparison (Corr), with the density of data points indicated on the diagonal plots.

**Figure S3.**
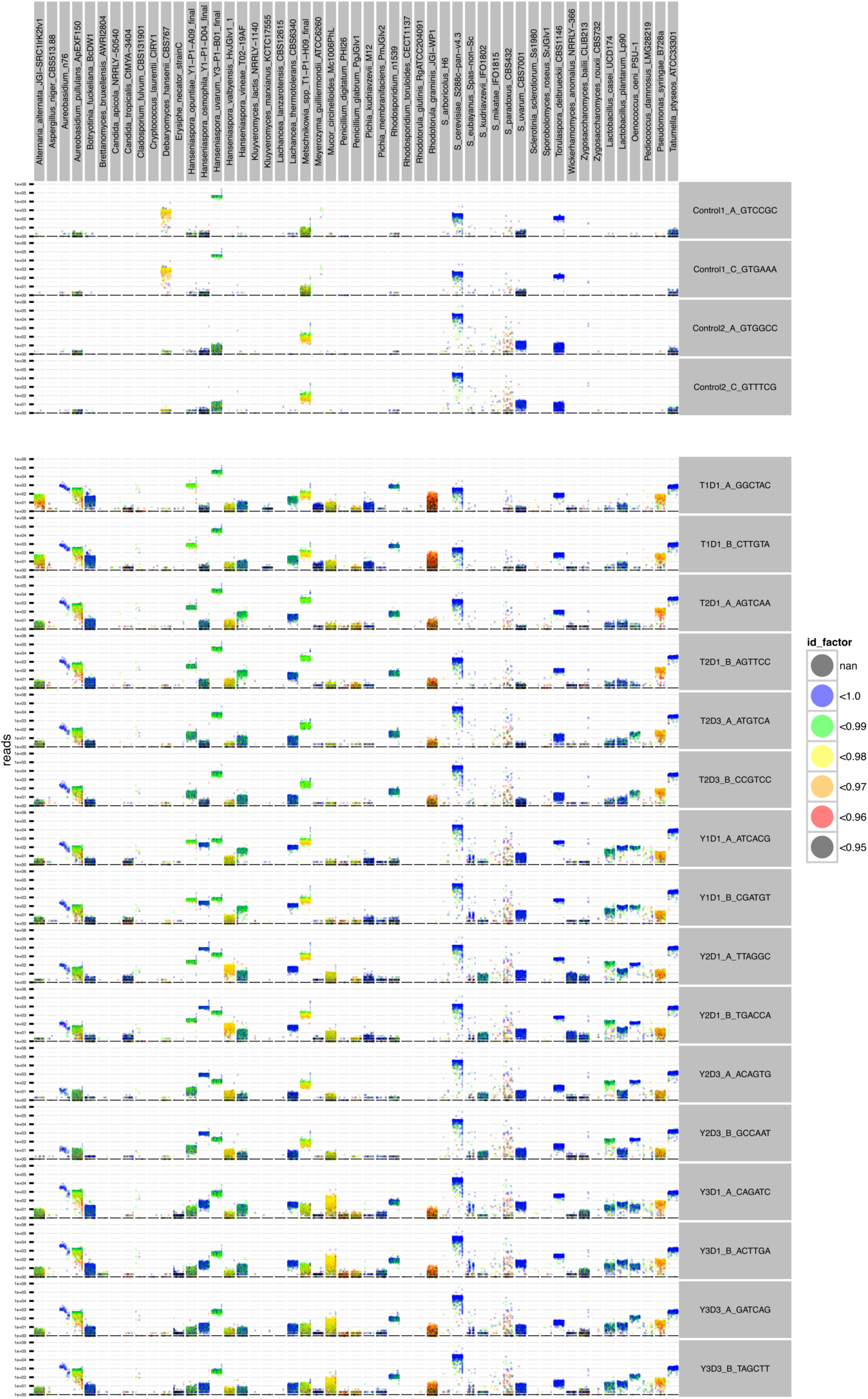
Shotgun metagenomic analysis of species abundance in wild fermentation via read mapping. Shotgun sequencing reads from each sample were mapped to the wine metagenome reference set. The total reads present in non-overlapping 10 kb windows across each genome were recorded relative to genomic location. In addition to total read number, the average identity of the reads in each window compared to the reference sequence was also calculated (id_factor).

